# On the heritability of criminal justice processing

**DOI:** 10.1101/107748

**Authors:** Brian B. Boutwell, Eric Connolly

## Abstract

An impressive number of researchers have devoted a great amount of effort toward examining various predictors of criminal justice processing outcomes. Indeed, a vast amount of research has examined various individual- and aggregate-level predictors of arrests, incarceration, and sentencing decisions. To this point, less attention has been devoted toward uncovering the relative contribution of genetic and environmental effects on variation in risk for criminal justice processing. As a result, the current study employs a behavioral genetic design in order to help fill this void in the existing literature. Using twin data from a national sample of youth, the current study produced evidence suggesting that genetic factors accounted for at least a portion of variance in risk for incarceration among female twins and probation among male twins. Shared and nonshared environmental influences accounted for the variance in risk for arrest among both female and male twins, probation among female twins, and incarceration among male twins. Ultimately, it appears that risk for contact with the criminal justice system and criminal justice processing is structured by a combination of factors beyond shared cultural and neighborhood environments, and appear to also include genetic factors as well. Moving forward, continuing to not use genetically sensitive research designs capable of estimating the role of genetic and nonshared environmental influences on criminal justice outcomes may result in misleading results.

---

One of the overarching questions in criminological and criminal justice research concerns uncovering variables that predict contact with the criminal justice (CJ) system (Sampson and Lauritsen, 1997; Owen, 2014). To be sure, a wealth of research has examined factors thought to increase the likelihood of arrest and formal processing and little debate exists concerning at least one fact: contact with the CJ system is not a random occurrence (Sampson and Lauritsen, 1997). Indeed, demographic factors including race (Sampson and Lauritsen, 1997; Walsh, 2004), age (Moffitt, 1993), and gender (Ferguson and Horwood, 2002), as well as personality traits and developmental processes (Gottfredson and Hirschi, 1990; Moffitt, 1993) all correlate with varying levels of risk for arrest, incarceration, and formal sentencing. What is noticeably lacking, however, is an effort to examine the role of genetic factors in predicting contact with the CJ system (Owen, 2014).

The evidence implicating genetic factors as source of variance for antisocial and criminal behavior in general is overwhelming (Ferguson, 2010; Mason and Frick, 1994; Miles and Carey, 1997; Plomin et al., 2013; Rhee and Waldman, 2002). The vast majority of researchers in criminology, however, have labored under the assumption that genetic factors play only a minimal part in predicting whom in the population will be arrested, incarcerated, and formally sentenced (Cullen, 2011). This approach may ultimately prove short sighted given the wide array of human outcomes influenced by genetic factors (Chabris et al., 2014; Turkheimer, 2000). The current study, thus, is intended to be a step toward uncovering sources of variation—both environmental and genetic—for CJ processing. First, however, it is important to discuss prior research pertaining to the heritability of antisocial, aggressive, and criminal behaviors.

## THE HERITABILITY OF ANTISOCIAL BEHAVIOR

Evidence concerning the heritability of human behavior can be traced largely to work in the field of behavior genetics (Plomin et al., 2013). The science of behavior genetics is devoted to examining the origin of individual differences (Turkheimer, 2000). Put differently, behavioral genetic research utilizes sibling pairs of varying degrees of genetic relatedness (i.e., monozygotic (MZ) twins, dizygotic (DZ) twins, full-siblings, and half-siblings) in order to uncover sources of variation for physiological, pathological, psychopathological, and behavioral outcomes (Turkheimer, 2000). Because MZ twins share, on average, twice as much genetic material as DZ twins, it is assumed that if MZ twins resemble one another more closely than DZ twins for a certain trait then genetic factors may be contributing to variation in that particular outcome (Plomin et al., 2013). If the similarity of DZ twins and full-siblings, which share on average 50% of their genetic material, is greater than that of half-siblings, which share 25% of their genetic material, moreover, the conclusion regarding genetic influences is further underscored (Plomin et al., 2013).

Put more directly, behavioral genetic research divides trait variance into that which is the result of heritability (*h*^2^), that which is the result of the shared environment (*c*^2^), and that which is due to the non-shared environment (*e*^2^) (Plomin et al., 2013). Prior researchers have discussed these concepts in great detail so they will not be belabored here (for a thorough methodological treatment of various concepts and assumptions of twin models see Barnes et al., 2014). For definitional purposes, however, heritability refers the proportion of variance in a given trait due to variation at the genetic level (Plomin et al., 2013). The shared environment represents environmental influences that serve to increase the resemblance of twins or siblings on the outcome measure under investigation. Non-shared environments represent the unique experiences of twins or siblings that function to create differences on the outcome measure under investigation between children in the same family. Taken together, *h*^2^, *c*^2^, *e*^2^ will always sum to yield 100 percent of the variance in a given outcome measure (Plomin et al., 2013).

The application of behavior genetic techniques is common in a range of disciplines outside of criminology (Rhee and Waldman, 2002). Behavior genetic methodologies, however, have recently gained some traction in the study of crime and delinquency. This growing line of research has produced evidence that abstention from delinquency (Barnes et al., 2011), onset of delinquency (DeLisi et al., 2008), chronic criminality (Barnes and Boutwell, 2012), changes in delinquency (Connolly, Schwartz, Nedelec, Beaver, and Barnes, 2015) delinquent peer affiliation (Beaver et al., 2009), and victimization (Barnes et al., 2011; Beaver et al., 2009) are all, to some degree, heritable.

Personality traits corresponding to risk of criminal and delinquent behavior have also shown evidence of being under genetic influence (Krueger et al., 2008; Rhee and Waldman, 2002). Studies, for example, examining low self-control (Beaver, Ratchford, and Ferguson, 2009; Beaver et al., 2008; Boisvert et al., 2011; Connolly and Beaver, 2014; Wright and Beaver, 2005; Wright et al., 2008), negative emotionality (Krueger et al., 2008), and psychopathy (Beaver et al., 2011) have produced consistent evidence that personality constructs positively correlated with crime and criminality are influenced by genetic factors. Equally important, environmental exposures, in general, do not occur randomly and seem to be at least partly heritable (Kendler and Baker, 2007). Taken as a whole, what this body of evidence suggests is that the behaviors that are likely to increase the risk of arrest (i.e., delinquency), as well as the personality constructs that are associated with those behaviors (i.e., self-control), are influenced to varying degrees by genetic factors. What is less clear, however, is whether actual instances of contact with the CJ system—including arrests and convictions—are heritable. Despite limited research in this regard, there is reason to suspect that formal processing through the CJ system may be influenced by genetic factors (for a broad overview of a related topic, see Kendler and Baker, 2007).

## GENETIC CONTRIBUTORS OF CRIMINAL JUSTICE PROCESSING

Early evidence concerning the role of genes in predicting CJ processing outcomes was produced by Mednick and his colleagues (1984) using a sample of adoptees. Mednick et al. (1984) examined both property and violent criminal convictions in adopted away children and their biological parents. While the results indicated that genetic factors may influence convictions for property crime (adopted children correlated significantly with their biological parents for this variable), there was no significant correlation between adopted children and biological parents for violent criminal convictions.

A more recent analysis of adoptees was conducted by Beaver (2011) using data on individuals participating in the National Longitudinal Study of Adolescent to Adult Health (Add Health). In order to indirectly tap genetic risk for antisocial behavior, subjects who reported having a biological parent previously incarcerated were considered to be genetically vulnerable for criminal involvement. The results were indeed stark. Subjects, for example, who reported two biological parents with a criminal history were over four times more likely to be arrested, over four times more likely to be incarcerated, over eight times more likely to be sentenced to probation, and approximately eight times more likely to be arrested multiple times.

Using a large sample based outside of the United States, Frisell, Lichtenstein, and Långström (2011) examined convictions for violent crime in a sample consisting of millions of Swedish residents. The analyses examined familial aggregation across a range of criminal convictions, including convictions for homicide, assault, robbery, and arson (as well as other types of illegal actions). The results provided compelling evidence of familial aggregation for multiple forms of criminal conviction. Importantly, not only did relatives resemble one another for criminal convictions, the similarity generally increased along with genetic relatedness of the family members (though mating couples also displayed similarity). This type of “genetic cascade” (Plomin et al., 2013) is suggestive of a heritable trait. And as Frisell et al. (2011) correctly point out, their results map with those of prior behavioral genetic studies revealing that at least a moderate proportion of the variance in criminality is the result of genetic variation (Rhee and Waldman, 2002). Lacking in this study, however, were precise parameter estimates for the contribution of genetic and environmental factors to trait variance.

## THE CURRENT STUDY

The studies just described provide an indication that CJ processing may be influenced by genetic factors, however, there is less evidence directly pertaining to the heritability of contact with the CJ system. Thus, the current study is intended to directly test this proposition. It is important to note that the current study utilizes the same dataset and CJ processing variables as prior research (Beaver, 2011; Beaver and Chaviano, 2011; Schwartz and Beaver, 2011). This analysis, however, differs from earlier work in at least four important ways. First, Beaver (2011) utilized adopted siblings in the Add Health and the current study uses the subsample of biologically related siblings. The current analysis excludes any siblings who were unrelated, yet residing in the same home. Essentially, the sample of participants analyzed here represents a different sample from that of Beaver (2011).^1^

Second, Beaver’s (2011) analysis employed an indirect measure of genetic risk to estimate the broad effects of the genome on risk for criminal justice processing. The current study analyzes sibling pairs in order to calculate pair similarity for a range of important CJ processing variables. Third, using behavioral genetic techniques, the current study is intended to estimate the proportion of variance in CJ processing that is accounted for by latent genetic and environmental influences. This was not the intent of prior research (Beaver, 2011). Fourth, and finally, prior researchers examining the role of genetic factors in CJ processing have examined the impact of measured genes (Beaver & Chaviano, 2011; Schwartz & Beaver, 2011). Researchers have become increasingly aware, however, that candidate gene research has some inherent limitations (Chabris et al., 2015; Dick et al., 2015). As such, tremendous caution should be taken when examining the impact of individual genes on complex behaviors.

## METHODS

### Data

The data included in this study were taken from the fourth wave of the National Longitudinal Study of Adolescent to Adult Health (Add Health) (Harris et al., 2006). The Add Health is a multi-wave sample spanning decades of development beginning in early adolescence and traversing into adulthood. Data collection began in 1994 while respondents were enrolled in middle and high school. Three waves of subsequent data collection were undertaken with respondents at each wave reporting on topics ranging from their personality traits, sexual activity, criminal involvement, victimization, family structures and existing medical conditions.

A useful feature of the Add Health sample is that sibling pairs, of varying degrees of genetic relatedness (i.e., monozygotic (MZ) twins, dizygotic (DZ) twins, full siblings, half-siblings, and non-related siblings (i.e., step-siblings living together in the same home), were actively recruited for participation in data collection (Harris et al., 2006). Subjects indicating that they had a sibling were (along with their co-twin) selected with 100 percent certainty (Harris et al., 2006). The resulting sampling procedure netted 307 MZ twin pairs and 452 DZ twin pairs. The current study analyzes twin pairs who provided a response to each CJ outcome. As such, the final analytic sample included 214 MZ twin pairs and 326 DZ twin pairs.

### Measures

*CJ Contact*. During the fourth, and most recent, wave of data collection respondents in the Add Health sample were asked to indicate whether they had ever been arrested, incarcerated, or placed on probation (Beaver, 2011; Beaver & Chiavano, 2011; Schwartz & Beaver, 2011). Responses to the arrest and incarceration items were coded dichotomously such that 1 = yes and 0 = no. Responses to the item assessing probation, however, was originally coded so that 0 = zero times, 1 = once, and 2 = more than once. For ease of interpretation and consistency with other CJ processing measures, the item assessing probation during the fourth wave was dichotomized such that 1 = ever convicted/placed on probation and 0 = no convictions/probation sentences.

## PLAN OF ANALYSIS

The plan of analysis for the current study involves a series of steps intended to systematically examine the genetic and environmental contributions to contact with the CJ system and CJ processing. The first step involved calculating between-sibling correlations for MZ twins and DZ twins to estimate concordance rates for arrest, incarceration, and probation. This step was carried out to examine whether genetic influences may explain a degree of variation in liability for each outcome. If concordance on an outcome for MZ twins (who share close to 100% of their genetic material) is stronger than concordance for DZ twins (who share, on average, 50% of their genetic material), then this can be interpreted as preliminary evidence suggesting that genetic factors account for a degree of variation in the outcome measure.

The second step was to calculate intraclass odds ratios for each of the outcomes described above (Cho et al., 2006). The logistic regression equation estimated in this phase of the analytical process assumes the following form:

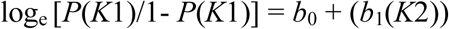

The interpretation of coefficient *b*_1_ remains straightforward, although it departs slightly from other forms of logistic regression analysis. In this case, the coefficient can be transformed (i.e., exponentiated in order to represent an odds ratio) so that the parameter estimate captures the odds that sibling 1 has a 1 on the outcome variable based on their sibling’s (i.e., sibling 2) score on the same measure—in this case one of the criminal justice processing items (Barnes et al., 2011). Importantly, genetic influences from this portion of the analysis are inferred if the similarity of sibling pairs increases along with their genetic relatedness (i.e., if MZ twins are more similar than DZ twins) (Plomin et al., 2013).

The third step in study was to directly estimate the proportion of variance in each criminal justice measure as well as the composite criminal justice measure that was the result of genetic and environmental influences. In order to estimate these effects, a series of univariate ACE models were fitted to the data. Biometric ACE models are capable of partitioning variance of a measurable outcome into three latent variance components: an additive genetic component (symbolized as A), a shared environmental component (symbolized as C), and a nonshared environmental component, which also includes measurement error (symbolized as E). Variation in an outcome explained by additive genetic effects suggests that genetic differences between-sibling pairs explain individual differences in the outcome under investigation.

Variation in an outcome explained by shared environmental effects suggests that shared environmental experiences between siblings explain similarities in a given outcome under investigation. In contrast, variation in an outcome accounted for by nonshared environmental effects suggests that unique environmental experiences between siblings explain differences in an outcome under investigation. The nonshared environmental component also includes the effects of measurement error. The assumptions of the twin-based research design have been tested several different times and in many different ways with results indicating that the twin-based research method produces reliable and stable estimates of genetic and environmental effects on phenotypic variance, even when underlying assumptions are violated (Barnes et al., 2014).

Given the dichotomous nature of the data, liability-threshold ACE models were estimated to assess the magnitude of additive genetic, shared environmental, and nonshared environmental effects on variation in liability for arrest, incarceration, and probation (Prescott, 2004). ACE models were estimated using the statistical software program M*plus* 7.1 (Muthén & Muthén, 1998-2011) with a weighted least squares estimator (i.e., WLSMV). Model fit was evaluated based on an adjusted χ^2^ difference test statistic, the comparative fit index (CFI), and the root mean square error of approximation (RMSEA). Based on prior literature, the following model fit cutoff points were used to assess satisfactory model fit: CFI ≥ .90, TLI ≥ .95, and RMSEA ≤ .05 (Hu & Bentler, 1999).

## RESULTS

Table 1 presents the descriptive statistics for each outcome measure. As can be seen, close to 28% of the twin sample reported having been arrested in their lifetime. Males were significantly more likely to report having been arrested compared to females ([1], χ^2^ = 44.96, *p* < .01). Table 1 also shows that 15% of the twin sample reported having been incarcerated in their lifetime and close to 12% of the sample reported having been on probation in their lifetime. Males were significantly more likely to report having been incarceration ([1], χ^2^ = 36.20, *p* < .01) and on probation ([1], χ^2^ = 54.67, *p* < .01) compared to females.

**Table 1.** Descriptive Statistics

| | Percent | Min | Max | *N* | (*df*) $\chi^2$ |
| --- | --- | --- | --- | --- | --- |
| Arrest |  |  |  |  |  |
| Total Sample | 27.88% | 0 | 1 | 1,090 | (1) 44.96** |
| Female | 18.99% | 0 | 1 | 558 |  |
| Male | 37.21% | 0 | 1 | 532 |  |
| Incarceration |  |  |  |  |  |
| Total Sample | 15.59% | 0 | 1 | 1,090 | (1) 36.20** |
| Female | 9.13% | 0 | 1 | 558 |  |
| Male | 22.36% | 0 | 1 | 532 |  |
| Probation |  |  |  |  |  |
| Total Sample | 11.92% | 0 | 1 | 1,090 | (1) 54.67** |
| Female | 4.83% | 0 | 1 | 558 |  |
| Male | 19.36% | 0 | 1 | 532 |  |

Notes: \*\* $p < .01$

Table 2 contains the between-sibling tetrachoric correlation estimates for arrest, incarceration, and probation. Estimates from the full twin sample including same-sex female and male twins revealed that MZ twins reported stronger concordance for arrest, incarceration, and probation, compared to DZ twins. However, the pattern of correlations was slightly altered when examining female and male twins separately. Specifically, concordance rates for arrest were slightly stronger for female MZ twins compared to same-sex female DZ twins and considerably stronger for incarceration. However, same-sex female DZ twins reported slightly stronger rates of concordance for probation compared to female MZ twins, but the correlations were nonsignificant. With respect to male twins, male MZ twins demonstrated slightly stronger concordance for arrest compared to same-sex male DZ twins and moderately stronger rates of concordance for probation. The strength of with-pair concordance for incarceration between male MZ and same-sex DZ twins was almost identical and non-significant.

**Table 2.** Twin Correlations for Arrest, Incarceration, and Probation

|  | Tetrachoric Correlations |  |  |
| --- | --- | --- | --- |
|  | Arrest | Incarceration | Probation |
| Total Sample |  |  |  |
| MZ | .62** | .44** | .53** |
| DZ | .48** | .28* | .38** |
| Female |  |  |  |
| MZ | .59** | .59** | .45 |
| DZ | .45** | .31* | .50 |
| Male |  |  |  |
| MZ | .63** | .26 | .50* |
| DZ | .51** | .25 | .35* |

Notes: \*\* $p < .01$ ; \* $p < .05$

Table 3 contains the results of the intraclass logistic regression analysis described in the plan of analysis. As can be seen in Table 3, the odds of arrest were significantly higher if MZ twins reported having a co-twin who was also arrested (*OR* = 7.02, *p* < .01, 95% CI: 3.48-14.18). This estimate was higher compared to DZ twins who reported having a co-twin who was also arrested (*OR* = 3.98, *p* < .01. 95% CI: 2.41-6.59). This pattern of results was consistent across all other outcomes when examining the full twin sample where the odds of incarceration and probation diminished as the degree of genetic relatedness decreased. When female and male twins were examined separately, however, the odds of arrest were significantly higher for female and male MZ twins compared to same-sex DZ twins, but the odds of probation for female MZ twins were non-significant and smaller compared to same-sex female DZ twins, while the odds of incarceration for male MZ and same-sex DZ twins were non-significant.

**Table 3.** Intraclass Odds Ratios for Arrest, Incarceration, and Probation

|  | Intraclass Odds Ratio |  |  |
| --- | --- | --- | --- |
|  | Arrest | Incarceration | Probation |
| Total Sample |  |  |  |
| MZ | 7.02**<br>[3.48-14.18] | 4.31**<br>[1.76-10.56] | 7.28**<br>[2.32-22.78] |
| DZ | 3.98**<br>[2.41-6.59] | 2.39*<br>[1.22-4.69] | 3.47**<br>[1.67-7.19] |
| Female |  |  |  |
| MZ | 6.58**<br>[2.41-17.97] | 7.88**<br>[2.21-28.10] | 7.06<br>[.61-42.71] |
| DZ | 4.08**<br>[1.62-10.25] | 3.12*<br>[1.63-13.25] | 8.12*<br>[1.06-57.51] |
| Male |  |  |  |
| MZ | 7.26**<br>[2.69-19.58] | 2.40<br>[.64-8.90] | 5.42*<br>[1.44-16.38] |
| DZ | 4.33**<br>[2.22-8.47] | 2.12<br>[.95-4.75] | 2.92*<br>[1.26-6.76] |

Notes: \*\* $p < .01$ ; \* $p < .05$

The final step in the analytic plan was to directly estimate the proportion of variance in criminal justice processing measures that was attributable to genetic and environmental effects. We fitted a series of univariate ACE models to the data on arrest, incarceration, and probation. Based on the evidence from previous steps in the analysis on sex differences in the magnitude of genetic and environmental effects on variation in risk for criminal justice processing, model fit statistics were used to evaluate whether model parameters could be equated across sex without significant loss of fit to the model. Results indicated that model parameters could be equated across sex without significant loss of fit for arrest ([3] Δχ^2^ = 3.51, *p* = .26), but not for incarceration ([3] Δχ^2^ = 1.03, *p* = .001) and probation ([3] Δχ^2^ = .09, *p* = .001). As such, a univariate ACE model was estimated to assess the magnitude of genetic and environmental effects on risk for arrest using the full twin sample, while ACE models for incarceration and probation were estimated separately for female and male twins. Table 4 presents the parameter estimates from a series of models examining the proportion of variance in risk for arrest accounted for by additive genetic, shared environmental, and nonshared environmental effects. As can be seen, constraining the additive genetic component to 0, which resulted in a CE model, improved overall model fit and provided the best fit to the data (Δχ^2^ = 5.77, Δdf = 1, *p* = .02, CFI = .97, RMSEA = .05). Standardized parameter estimates from the best-fitting CE model suggested that 54% of the variation in liability for arrest was accounted for by shared environmental effects, while 46% of the variation in liability was accounted for by nonshared environmental effects.

**Table 4.** Univariate ACE Estimates for Arrest for Female and Male Twins (*N* = 540 pairs)

| | | A | C | E | $\Delta\chi^2$ | $\Delta$ df | *p* | CFI | RMSEA |
| --- | --- | --- | --- | --- | --- | --- | --- | --- | --- |
| Models |  |  |  |  |  |  |  |  |  |
| Arrest | ACE | .28<br>[-.10-.66] | .34<br>[.04-.63] | .37**<br>[.23-.52] | - | - | - | .97 | .05 |
|  | AE | .70**<br>[.57-.83] | .00<br>[.00-.00] | .30**<br>[.17-.42] | 14.65** | 1 | <.001 | .93 | .07 |
|  | **CE** | **.00**<br>**[.00-.00]** | **.54\*\***<br>**[.44-.64]** | **.46\*\***<br>**[.36-.55]** | **5.77\*** | **1** | **.02** | **.97** | **.05** |
|  | E | .00<br>[.00-.00] | .00<br>[.00-.00] | 1.00***<br>[1.00-1.00] | 355.74** | 2 | <.001 | .00 | .26 |

*Notes:* Best-fitting ACE model is bolded. Results are standardized parameter estimates with 95% confidence intervals in parentheses.
\*\* $p < .01$ ; \* $p < .05$

Table 5 presents the standardized parameter estimates from the estimated ACE models examining the magnitude of genetic and environmental effects on risk for incarceration and probation among female twins. Model fit indices indicated that an AE model for incarceration fit the data best (Δχ^2^ = .03, Δdf = 1, *p* = .85, CFI = .93, RMSEA = .04) where additive genetic influences accounted for 60% of the variation in liability for incarceration and nonshared environmental influences accounted for 40% of the variation in liability. The best-fitting univariate model for probation included only shared and nonshared environmental parameters (Δχ^2^ = .00, Δdf = 1, *p* = 1.00, CFI = .80, RMSEA = .06). As a result, the parameter estimates from the best-fitting model suggested that 49% of the variation in liability for probation was accounted for by shared environmental influences, while 51% of the variance in liability was accounted for by nonshared environmental influences.

**Table 5.** ACE Parameter Estimates for Same-Sex Female Twins (*N* = 260 pairs)

| | A | C | E | $\Delta\chi^2$ | $\Delta$ df | $p$ | CFI | RMSEA |
| --- | --- | --- | --- | --- | --- | --- | --- | --- |
| Models |  |  |  |  |  |  |  |  |
| Incarceration |  |  |  |  |  |  |  |  |
| ACE | .55<br>[-.30-1.41] | .04<br>[-.69-.77] | .40*<br>[.14-.66] | - | - | - | .85 | .07 |
| **AE** | **.60\*\***<br>**[.35-.84]** | **.00**<br>**[.00-.00]** | **.40\*\***<br>**[.15-.64]** | **.03** | **1** | **.85** | **.93** | **.04** |
| CE | .00<br>[.00-.00] | .49**<br>[.28-.70] | .51**<br>[.30-.71] | 16.48* | 1 | .03 | .85 | .06 |
| E | .00<br>[.00-.00] | .00<br>[.00-.00] | 1.00**<br>[1.00-1.00] | 65.91** | 2 | <.001 | .00 | .16 |
| Probation |  |  |  |  |  |  |  |  |
| ACE | .00<br>[.00-.00] | .49**<br>[.18-.79] | .51**<br>[.20-.81] | - | - | - | .61 | .10 |
| AE | .62*<br>[.20-1.04] | .00<br>[.00-.00] | .37<br>[-.04-.79] | 28.11** | 1 | <.001 | .72 | .08 |
| **CE** | **.00**<br>**[.00-.00]** | **.49\*\***<br>**[.18-.79]** | **.51\*\***<br>**[.20-.81]** | **.00** | **1** | **1.00** | **.80** | **.06** |
| E | .00<br>[.00-.00] | .00<br>[.00-.00] | 1.00**<br>[1.00-1.00] | 28.11** | 2 | <.001 | .00 | .13 |

Notes: Best-fitting ACE model is bolded. Results are standardized parameter estimates with 95% confidence intervals in parentheses.
\*\* $p < .01$ ; \* $p < .05$

Table 6 presents the standardized parameter estimates from the univariate ACE models for incarceration and probation among male twins. Model fit indices indicated that a CE model fit the data adequately (Δχ^2^ = .02, Δdf = 1, *p* = .88, CFI = .87, RMSEA = .09). Results from the CE model suggested that 26% of the variance in liability for incarceration among male twins was accounted for by shared environmental influences, while 74% of the variance in liability for incarceration was accounted for by nonshared environmental influences. With respect to probation, model fit statistics suggested that an AE model provided the best fit to the data ( Δχ^2^ = .1.44, Δdf = 1, *p* = .22, CFI = .90, RMSEA = .05). Parameter estimates from the best-fitting AE model indicated that 57% of the variance in liability for probation was accounted for by additive genetic influences, while the remaining 43% of variance in liability for probation was accounted for by nonshared environmental influences.

**Table 6.** ACE Parameter Estimates for Same-Sex Male Twins (*N* = 280 pairs)

| | A | C | E | $\Delta\chi^2$ | $\Delta$ df | p | CFI | RMSEA |
| --- | --- | --- | --- | --- | --- | --- | --- | --- |
| Models |  |  |  |  |  |  |  |  |
| Incarceration |  |  |  |  |  |  |  |  |
| ACE | .03<br>[-.78-.86] | .24<br>[-.33-.80] | .73<br>[.37-1.07] | - | - | - | .69 | .13 |
| AE | .36*<br>[.08-.63] | .00<br>[.00-.00] | .64**<br>[.36-.91] | 1.88 | 1 | .17 | .74 | .11 |
| **CE** | **.00**<br>**[.00-.00]** | **.26\***<br>**[.07-.45]** | **.74\*\***<br>**[.55-.92]** | **.02** | **1** | **.88** | **.87** | **.09** |
| E | .00<br>[.00-.00] | .00<br>[.00-.00] | 1.00***<br>[1.00-1.00] | 20.76** | 2 | <.001 | .00 | .15 |
| Probation |  |  |  |  |  |  |  |  |
| ACE | .31<br>[-.44-1.06] | .20<br>[-.34-.73] | .49**<br>[.19-.79] | - | - | - | .85 | .07 |
| **AE** | **.57\*\***<br>**[.31-.81]** | **.00**<br>**[.00-.00]** | **.43\*\***<br>**[.18-.68]** | **1.44** | **1** | **.22** | **.90** | **.05** |
| CE | .00<br>[.00-.00] | .40**<br>[.22-.58] | .60**<br>[.41-.77] | 1.84 | 1 | .17 | .88 | .06 |
| E | .00<br>[.00-.00] | .00<br>[.00-.00] | 1.00***<br>[1.00-1.00] | 57.49** | 2 | <.001 | .00 | .14 |

Notes: Best-fitting ACE model is bolded. Results are standardized parameter estimates with 95% confidence intervals in parentheses.
\*\* $p < .01$ ; \* $p < .05$

## DISCUSSION

Criminology has devoted a great deal of effort to understanding why some individuals in the population are more likely to come into contact the CJ system. Further understanding the underlying influences on the predictors of arrest, incarceration, and formal sentencing remain central to the research agendas for both criminologists and CJ practitioners. The current study was intended to press forward in this area of research by examining the genetic and specific environmental underpinnings of various measures of CJ processing (arrest, incarceration, and probation). While prior research suggests that CJ processing is not beyond the reach of genetic influence (Beaver, 2011; Beaver and Chaviano, 2011), less research has used behavioral genetic methods to assess the extent to which genetic and environmental influences account for variation in risk for CJ processing. The present study extended this line of work by examining a sample of male and female twins from a nationally representative survey of youth. The results yielded two key findings that deserve further attention.

First, our results revealed that shared and nonshared environmental influences accounted for the variance in liability for arrest among female and male twins. Interestingly, shared environmental influences explained the majority of the variance in risk for arrest suggesting that common environmental experiences play an important role in explaining why some come into contact with the CJ system. Although speculative, some possible mechanisms involved in the shared environmental influence on arrest might include shared subcultural values, socioeconomic factors, or even neighborhood factors (see also, Burt, 2009). Given the significant role of the non-shared environment as well, techniques such as those utilized by Beaver (2008) will be useful in further exploring whether differences in exposure to key crime correlates, like delinquent peer groups, might also be important.

Second, results from univariate biometric models examining female and male twins separately revealed that additive genetic and nonshared environmental influences explained variation in liability for incarceration among females and probation among males. Alternatively, the shared and nonshared environment appeared to explain variation in liability for probation among females and incarceration among males. While it is tempting to speculate about why genetic influences were different for different traits in males and females, we should resist making strong inferences until our findings can be replicating using larger, more powerful samples. Nonetheless, what is clear is that the non-shared environment consistently emerged as significant for every CJ processing variable. What this means is that future research should make use of twin and sibling designs in order to more thoroughly isolate and examine possible nonshared environmental causes and correlates of CJ processing (Beaver, 2008).

The current study was not without limitation and it is important to discuss potential shortcomings. The first limitation concerns the measurement utilized to assess formal contact with the criminal justice system. Respondents were asked to self-report their contact with the justice system, and as a result, issues with misreporting due to concerns over social desirability could potentially impact the findings reported here. There is some evidence suggesting that self-reported items are both reliable and valid instruments for assessing criminal involvement (Piquero, Farrington, and Blumstein, 2003; Thornberry and Krohn, 2000). The use of self-reported items, then, may help to side step many of the potential problems surrounding the use of official statistics (i.e., the justice system can only process crimes that are known to the police). Nonetheless, the extent to which self-reported items have a biasing effect on the results reported herein is an empirical question that requires further investigation. Also, while there is some evidence from the Add Health data that findings from the twin sample should generalize to a broader population of non-twins (Barnes and Boutwell, 2013), insight about the generalizability of CJ processing from twins to non-twins remains limited.

Ultimately, the findings presented here offer further evidence that various CJ outcomes are influenced by a combination of genetic, shared environmental, and nonshared environmental factors (Beaver, 2008; 2011; Frisell et al., 2011). When considered alongside all other evidence pertaining to the origins of virtually every other human outcome these results are to be expected (Chabris et al., 2014; Kendler and Baker, 2007; Turkheimer, 2000). Indeed, the sources of human variation for criminal behavior involve, to varying degrees, *both* genes and environments. Unfortunately, most criminological research examining CJ processing outcomes do not focus on estimating or controlling for latent genetic influences, which ultimately make it impossible to distinguish between shared and non-shared environmental effects (Beaver, 2008).

In light of the current results, future studies now have even more reason to employ genetically sensitive designs (Barnes et al., 2014). Such an assertion may seem odd, given that some of the best fitting models presented in the current study excluded the heritability variance component entirely, meaning that genetic influences did not appear to account for any of the variance in liability for CJ outcomes. This reality, however, does not obfuscate the importance of genetically *sensitive* research designs and the usefulness of twin data more broadly to criminological research. Importantly, commonly used standard social science approaches that do not account for the clustering of biologically related siblings in an analytic sample cannot disentangle shared from non-shared environmental influences (Barnes et al., 2014).

Though some of our models did reveal that genetic factors failed to account for any of variance in liability for CJ processing, every model revealed a significant non-shared environmental parameter. While interesting, it is important to remember that the nonshared environment component also contains variance from measurement error. That said, it remains critical for researchers to use genetically sensitive research designs such as the MZ difference score approach (Pike et al., 1996) or sibling comparison approach to further identify specific non-shared environments that might play an important role in explaining individual differences in CJ processing. Moreover, future research should begin to examine potential endophenotypes (e.g., intelligence, low self-control, psychopathy) involved in explaining sex-specific pathways between genetic influences and specific CJ processing outcomes. For the field of criminology to remain a relevant voice in the study of human behavior there needs to be a shift towards examining the relative contribution, and interplay, between genetic and environmental sources of variance on both criminal behavior and pathways to CJ processing (Barnes et al., 2014; Cullen, 2011).

## Footnotes

1 Beaver and Chaviano (2011) tested whether certain dopaminergic genes influenced the extent to which individuals were ever arrested, sentenced to probation, incarcerated, or arrested multiple times. For each outcome, genetic risk factors increased the likelihood of criminal justice processing. Schwartz and Beaver (2011) also employed a molecular genetics approach and tested whether certain environmental factors conditioned the influence of genetic factors in the prediction of criminal justice outcomes. Drawing on prior literature linking variants of the monoamine oxidase A (MAOA) gene to violent forms of criminal behavior (Beaver et al., 2009; Brunner et al., 1993). The findings from their study suggested that the impact of MAOA on being arrested was conditioned by the respondent’s perceived level of prejudice, dovetailing with work indicating that adverse environments condition the influence of MAOA in the prediction aggressive and violent behavior (Caspi et al., 2002; Kim-Cohen et al., 2006).

